# Active Site Local Environment Allows Acidic and Basic Synergy in Enzymatic Ester Hydrolysis by PETase

**DOI:** 10.64898/2026.02.03.703441

**Authors:** Jeffrey Fan, Xueqi Chu, Kevin Fang, Colin Zhang, Nicole Guo, Robert Chuang, Yujia Li, Jiaxu Chen, Emma Wang, Nina Ding, Joshua Franklin, Yang Ha

## Abstract

Polyethylene terephthalate (PET) is a commonly used plastic worldwide and reducing its prevalence is crucial to improving environmental pollution. PETase that degrades PET plastic have received a lot of attention recently. This paper evaluates the ester hydrolysis process under both acidic and basic conditions, and shows that the local environment of the protein active site takes advantage of both. High pH in the protein buffer creates a better nucleophile to attack the ester through a proton shuttle channel in the protein, while local hydrogen bonds to the carbonyl of the ester stabilizes the intermediate/transition state of the hydrolysis reaction. With the understanding at the atomic level, we propose two engineering directions that can potentially improve the reactivity of the PETase: 1) increase the alkaline stability of the protein in general; 2) perturb the local hydrogen bond network to increase the partial charge on the PET carbonyl to be hydrolyzed.

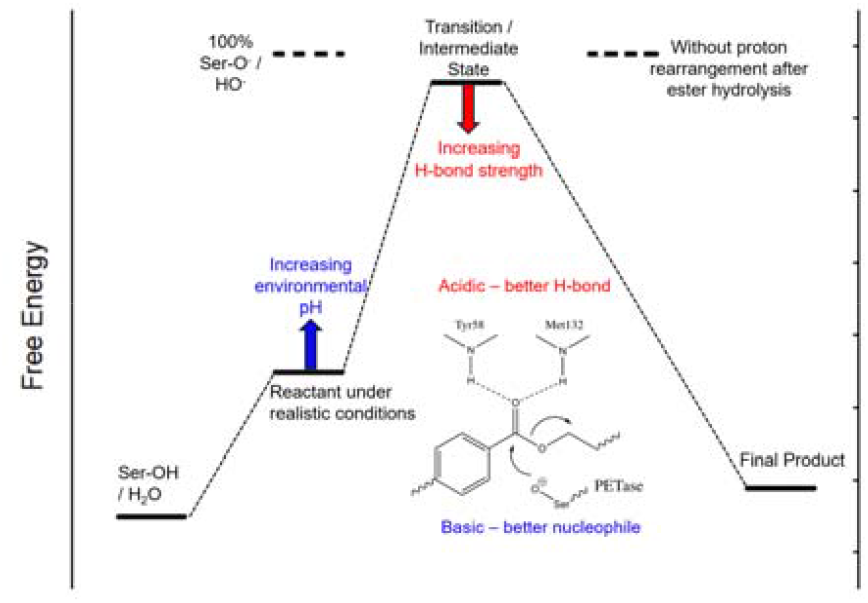

## 1 Introduction

Plastics are one of the most prevalent man-made materials globally. In the past 50 years, plastic production has increased 20-fold^1^, with most consisting of single-use packaging plastics^2^, which are disposed of rapidly and build up in landfills. Geyer et al.^2^ found that, in fact, of all the plastics in circulation up until 2015, only 9% have been recycled and are currently in circulation, whereas 79% have ended up in landfills or the environment. This buildup in the environment is due to plastics being non-biodegradable^3^, resulting in an ever-growing amount of plastic on the planet.

Plastics can break down over time through mechanical stresses, photo-oxidation, and biological processes^4^. However, the rate of degradation is not enough to recycle safe products back into the environment. The process of degradation creates secondary micro- and nano-plastics, with diverse chemical compositions, forms, and effects^5^. Primary microplastics are created through industrial processes directly and are often used in cosmetic products^5^ in the form of beads. The health risks posed by microplastics are non-negligible: toxic chemicals can be bound to these fibers or beads, including BPA, phthalates, polyfluorinated chemicals, and more. Microplastics have been found in the digestive tract, lungs^6^, circulation, reproductive systems^6^, and even the brain^7^, though the toxic^8^ effects of the mostly inert plastic particles have not been studied in much depth. As such, plastic pollution is a global health concern now.

Given the widespread prevalence of plastics in the environment, solutions must be proposed to reduce plastic pollution and leaching into the biosphere. We intend to target the first stage and ultimately degrade plastics into inert, smaller monomers through the biochemical degradation pathway. Our main focus is on polyethylene terephthalate (PET), one of the most widely used plastics for commercial and industrial applications. Compared with other plastics such as polyethylene (PE), which lack functional groups and are relatively inert, PET monomers are linked by ester bonds and are more prone to being degraded by enzymes. Numerous PETases have been discovered and studied extensively^9–12^. In one of the recent studies, Lu et al.^12^ used machine learning to aid enzyme engineering to improve the enzymatic ability through the modification of global protein stability. Other studies have improved the thermostability of IsPETase^13,14^. There are also theoretical studies, such as a QM/MM calculation to model the whole protein with the degradation process^15^. However, working with the whole protein tends to lack focus at the active site. Understanding the reaction mechanism at fundamental level may provide insights to best engineer the enzyme. In this work, we adopt a mechanism-informed strategy by focusing on the PETase active site and its reaction pathway. By evaluating the ester degradation reactions at the atomic scale and analyzing the energy profile, we have new insights into the advantage of the protein active sites and proposed directions to engineer PETase variants that surpass the efficiency of the native enzyme.

## 2 Materials and methods

Pymol^16^ was used to visualize the protein structure, and the PDB codes 5XJH and 5XH3 were used for detailed analysis. The reaction models with protein environments were built using Avogadro^17^. Density Functional Theory calculations were carried out using the ORCA software^18^. B3LYP functional and TZVP basis sets were chosen based on our previous experience, and a polarized continuum model (PCM) was used for solvent effects. The scan feature in ORCA was used to model the reaction coordinate and locate the transition state.

All calculations were run on the supercomputing cluster Lawrencium at the Lawrence Berkeley National Lab. We wrote our own Python scripts to pull out the energies from the output file and for further data analysis. Alphafold 3^19^ was used to check the fold of the protein after the proposed mutations.

## 3 Results and Discussion

### 3.1. Classic ester hydrolysis reactions

The nature of the PET degradation is essentially the hydrolysis of the ester bond, as shown in reaction (1). Organic chemistry textbooks say such reactions are not likely to occur under neutral conditions, but can occur under either acidic conditions or basic conditions. This is because under acidic conditions, the protonation of the carbonyl oxygen can better induce nucleophilic attack by water, while under basic conditions, the hydroxide ion itself is a better nucleophile than neutral water.

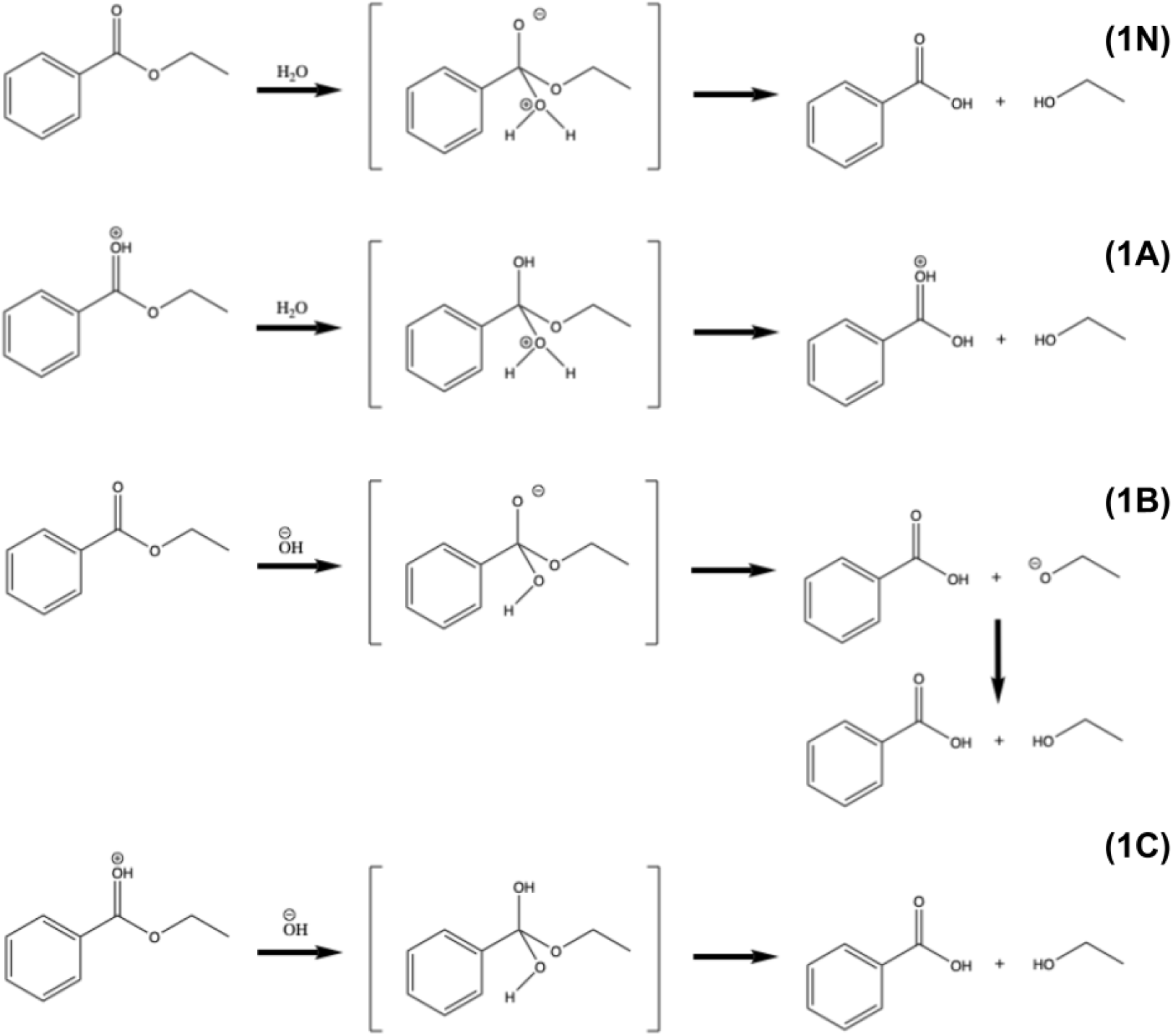

Computationally, the ΔE of the neutral reaction (1N) is +1.8 kcal/mol with a reaction barrier of 57.4 kcal/mol, suggesting that the reaction under neutral conditions is endothermic. The ΔE of the acidic reaction (1A) is 1.0 kcal/mol, slightly lower than that of neutral, but the reaction barrier decreases to 50.2 kcal/mol. On the other hand, the ΔE of the basic reaction (1B) is very exothermic, with -31.6 kcal/mol. A detailed investigation shows that the ΔE of the old C-O bond cleavage and new C-O bond formation is +2.0 kcal/mol, while the similar structure of the four-coordinated “transition state” becomes an intermediate state, which is 0.8kcal/mol lower than the reactant. The driving force is mainly from the later proton rearrangement from the carboxylic acid to the alcohol (Figure 1).

**Figure 1.**
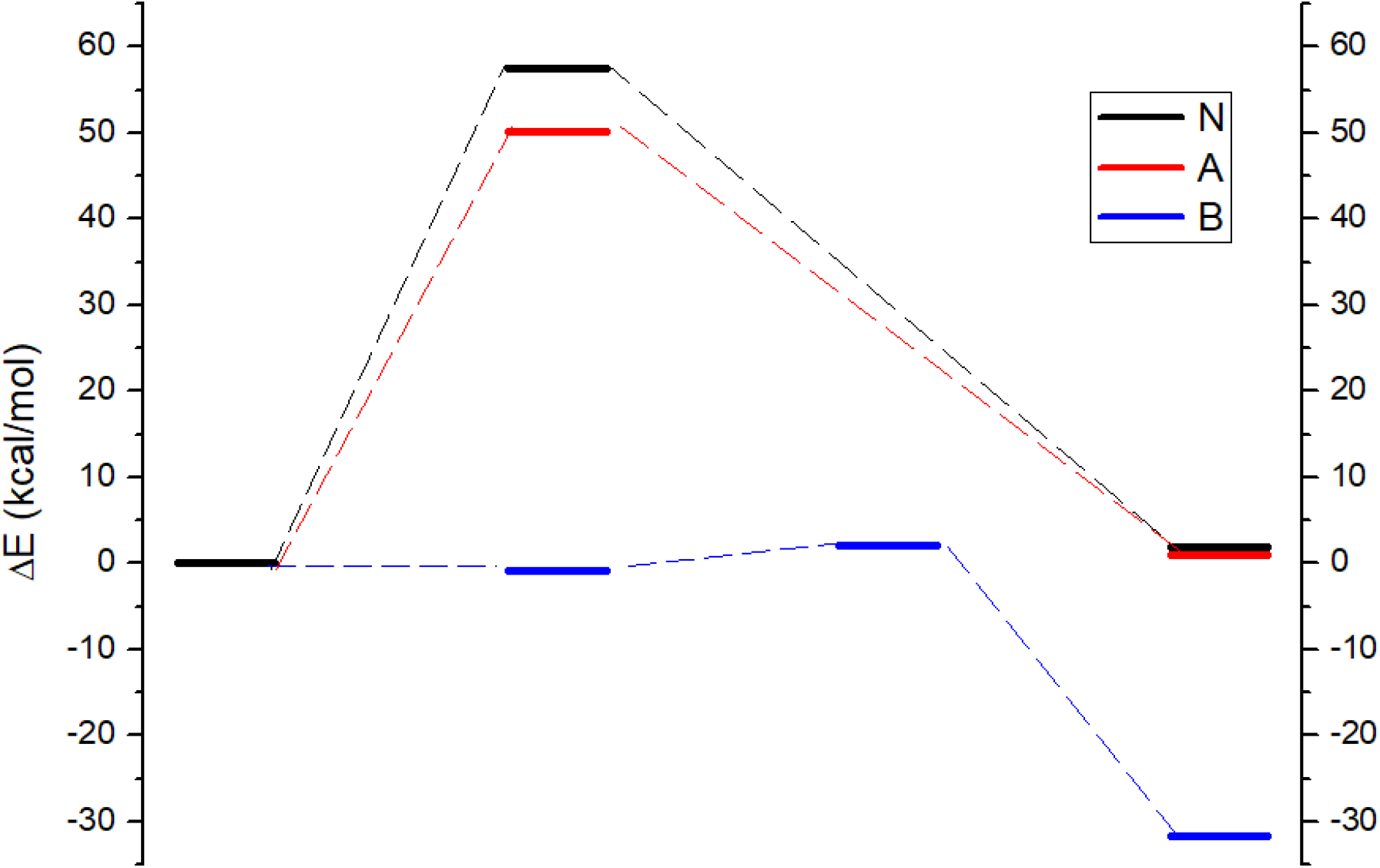
Energy profile for the ester hydrolysis under neutral (black), acidic (red), and basic (blue) conditions.

These calculated values are consistent with previous literature^20^ as well as the experimental observations that the hydrolysis of the ester is likely to occur under either acidic or basic conditions, and the reaction under basic conditions has more driving force. It should be noted that the calculations here, although using a PCM model for the solvent dielectric effect, are only applicable to molecular systems and cannot reflect the proton exchange in aqueous conditions. Thus, in reality, where the OH^-^ concentration is 10^-4^M for a pH=10 solution, the water concentration is 55.5M. The real free energy values should be much lower and will be discussed in later sections.

One can imagine that if the carbonyl is protonated and the nucleophile is a hydroxide ion, the reaction is more prone to occur. Calculations support this hypothesis that the calculated ΔE of reaction (1C) is even more negative to be -66.3kcal/mol. A similar set of calculations on ethyl acetate shows a comparable trend. Such mixed conditions are not possible in aqueous solution, but they can be possible in proteins, where the local reaction environment is determined by amino acids near the active site.

### 3.2. Ester hydrolysis in PETase

As shown in Figure 2, the active site of the PETase has a very conserved oxyanion hole. The carbonyl of the ester is hydrogen-bonded to two amides, which makes it partially protonated. On the other side, the deprotonated serine can be the nucleophile that readily attacks the carbonyl from the back. The first step is a transesterification (ester exchange) reaction, where the C-O bond in the PET is cleaved, and the carbonyl is connected to the protein serine, forming a new ester. After that step, another water or hydroxide enters the active site, cleaves the C-O bond between the left over PET and the serine, and regenerates the active site.

**Figure 2.**
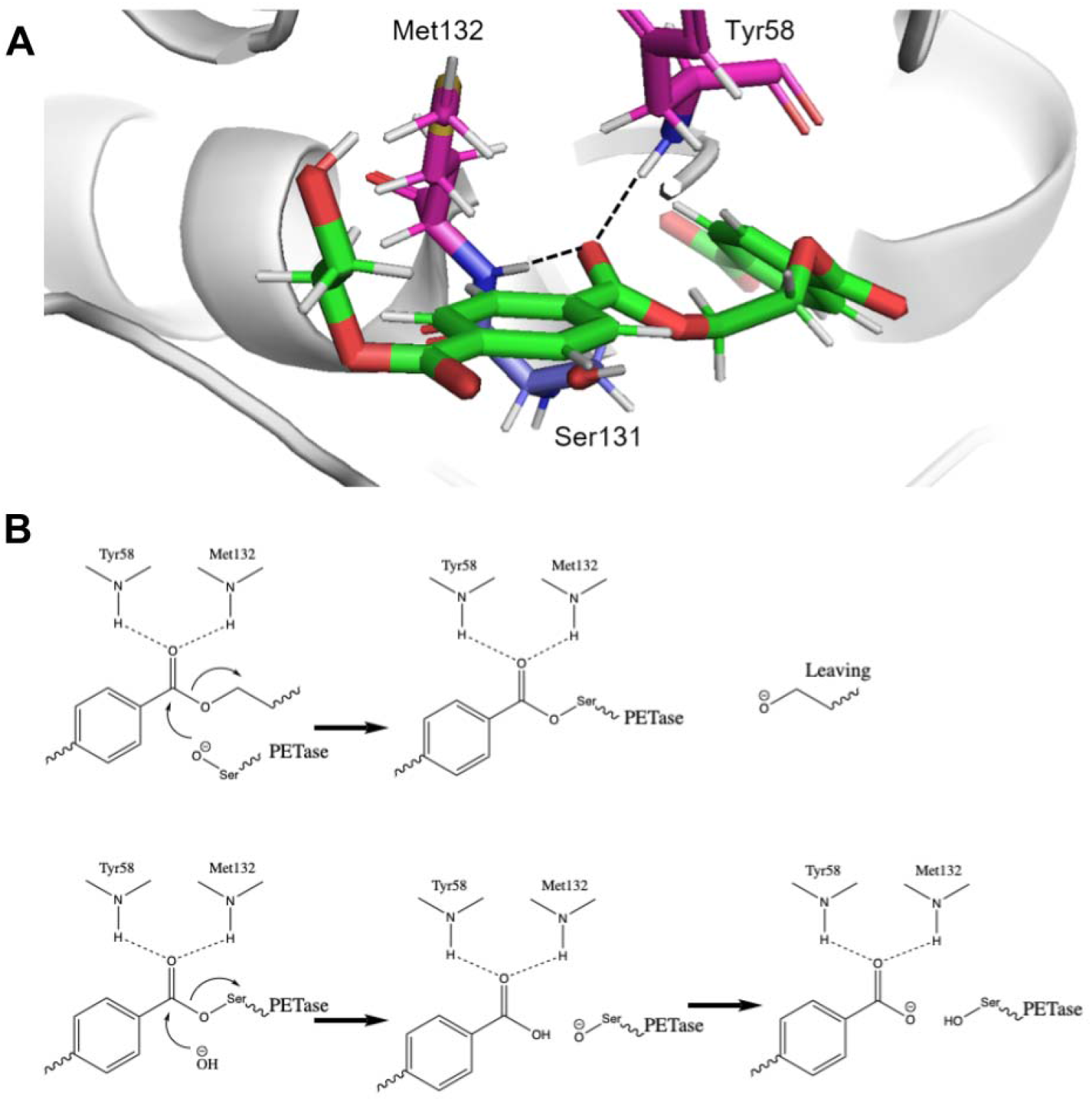
(A) The active site of the PETase, with the substrate shown in green, nucleophile serine shown in blue, and two amide hydrogen bond donors shown in magenta. (B) The two-step equations for the ester bond hydrolysis are shown in two steps. The first step is a transesterification (ester exchange) reaction, and the second step is the hydrolysis of the newly formed ester.

The overall reaction of the two steps will be similar to Rxn 1. Regardless of the role of the protein, the net thermodynamics of the ester hydrolysis cannot be changed. Based on the calculations above, such reactions are endothermic under neutral and acidic conditions. That means it requires energy input to achieve the PET degradation, thus the forward reaction is favored at relatively high temperatures. This is consistent with the fact that lots of the reactions in literature happen at up to 65 °C, and it explains the motivation in some studies for optimizing the thermostability of the enzyme^13, 14^.

However, based on the calculations in 3.1, there are other ways to potentially improve the enzyme performance. As shown above, the reaction is exothermic under basic conditions; thus, increasing the basicity of the reactants can make the reaction occur simultaneously at moderate temperatures. On the other hand, increasing the acidity around the carbonyl, although it cannot change the overall thermodynamics, can lower the reaction barrier for both ester exchange and the later hydrolysis step.

### 3.3. Improving the basicity of the nucleophile

The first ester exchange step can be initiated by deprotonation of the serine, while the following ester hydrolysis step is also favored if the nucleophile is a hydroxyl anion. Both anion formation reactions can occur through a proton shuttle mechanism by transferring the proton to the solvent environment (Figure 3).

**Figure 3.**
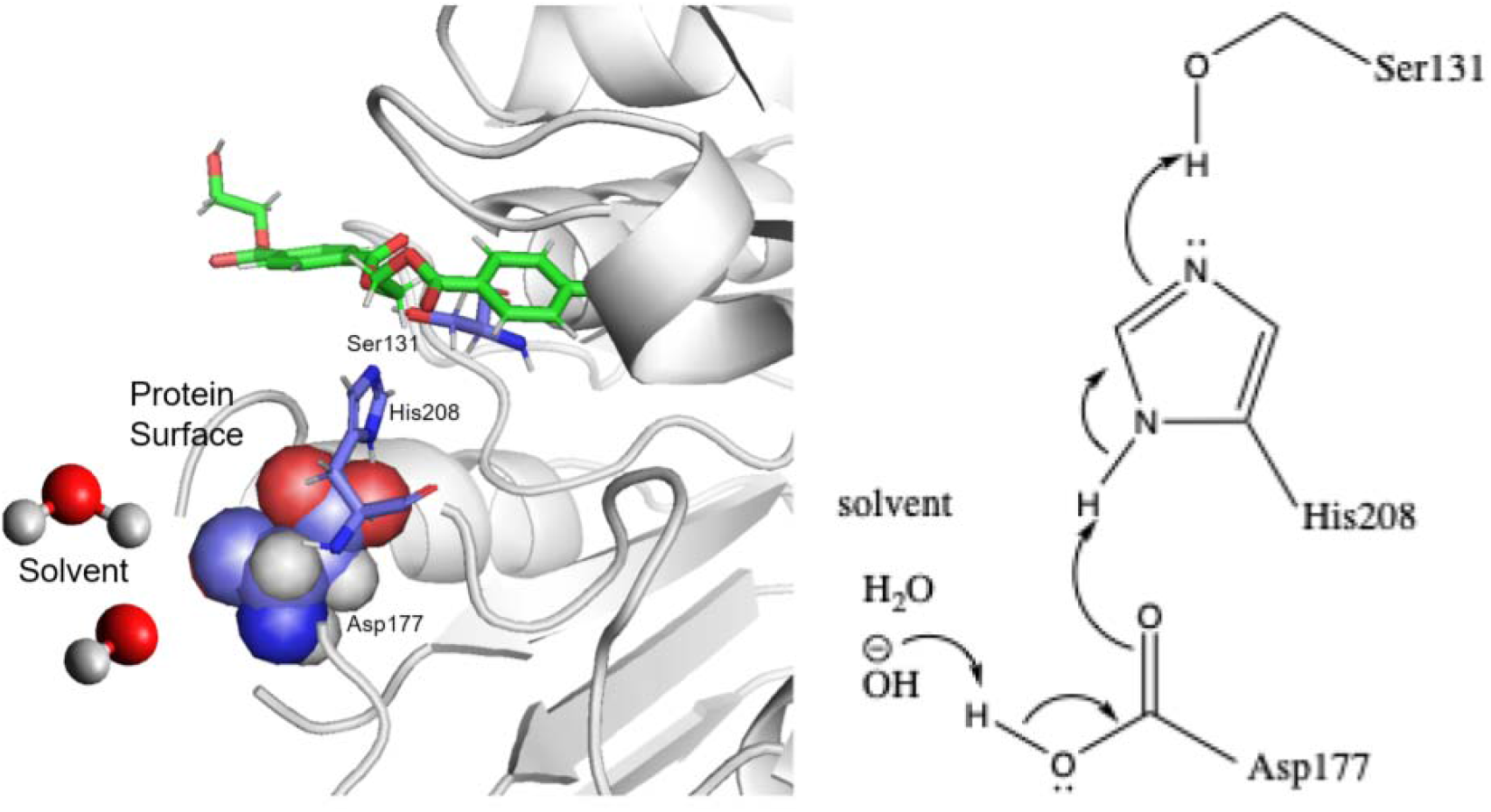
Demonstration that the pH of the solvent can impact the protonation state of Ser131 through a proton shuttle mechanism, which plays an important role in the ester hydrolysis.

Increasing the pH in the environment can increase the odds of the formation of the anionic nucleophile at the active site, leading to a more favorable forward reaction towards the ester hydrolysis. This is supported by the observation in Carletti et al.^21^. Improving the alkaline stability of the protein extends the lifetime of the enzyme, thus it can be a direction of protein engineering. Achieving alkaline stability, like enhancing thermostability, will require a holistic understanding of the entire protein — an important direction for future work.

### 3.4. Increasing the carbonyl protonation

As shown in 3.1, protonation of the carbonyl of the ester can effectively lower the reaction barrier of both the ester exchange and the final hydrolysis steps. In the enzyme, this is achieved by forming hydrogen bonds with two amides (Tyr58 and Met132) from the protein framework. We propose that increasing the H-bond strength can decrease the activation energy and validate this idea by DFT calculations. The experimental pKa values of amide H can be found in Šebesta et al.^22^, but it needs to be pointed out that modeling the pKa values of amide H by theory itself can be very challenging and worth a separate study.^22^ In this project, we did a series of 2D energy surface scans at different fixed H-bond distances (Figure 4), assuming longer distances lead to weaker H-bonds. This assumption was confirmed by the fact that the partial Mulliken charge on the carbonyl increases with shorter H-bonds. The relative energies of the transition/intermediate states at different hydrogen bond distances are shown in Table 1. It is clear that as the hydrogen bond becomes closer thus stronger, the transition/intermediate state is more stable, thus the reaction rate can go faster. This conclusion is also supported by the QM/MM calculations by Jerves et al.^15^, where they also reported the hydrogen bond shrinking at the transition state when they model the whole reaction coordinate.

**Table 1.** Transition/Intermediate State Energy Change Relative to H-bond Distance Between the Oxygen in Carbonyl in the PET and Amide Nitrogen in Tyr58 and Met132.

| N-H...O=C distance (Å) | Relative energy (kcal/mol) of the transition/intermediate state |
| --- | --- |
| No H-bond | 0 |
| 3.7 | -0.9 |
| 3.5 | -1.3 |
| 3.3 | -2.0 |
| 3.1 | -3.0 |
| 2.9 | -4.5 |

**Figure 4.**
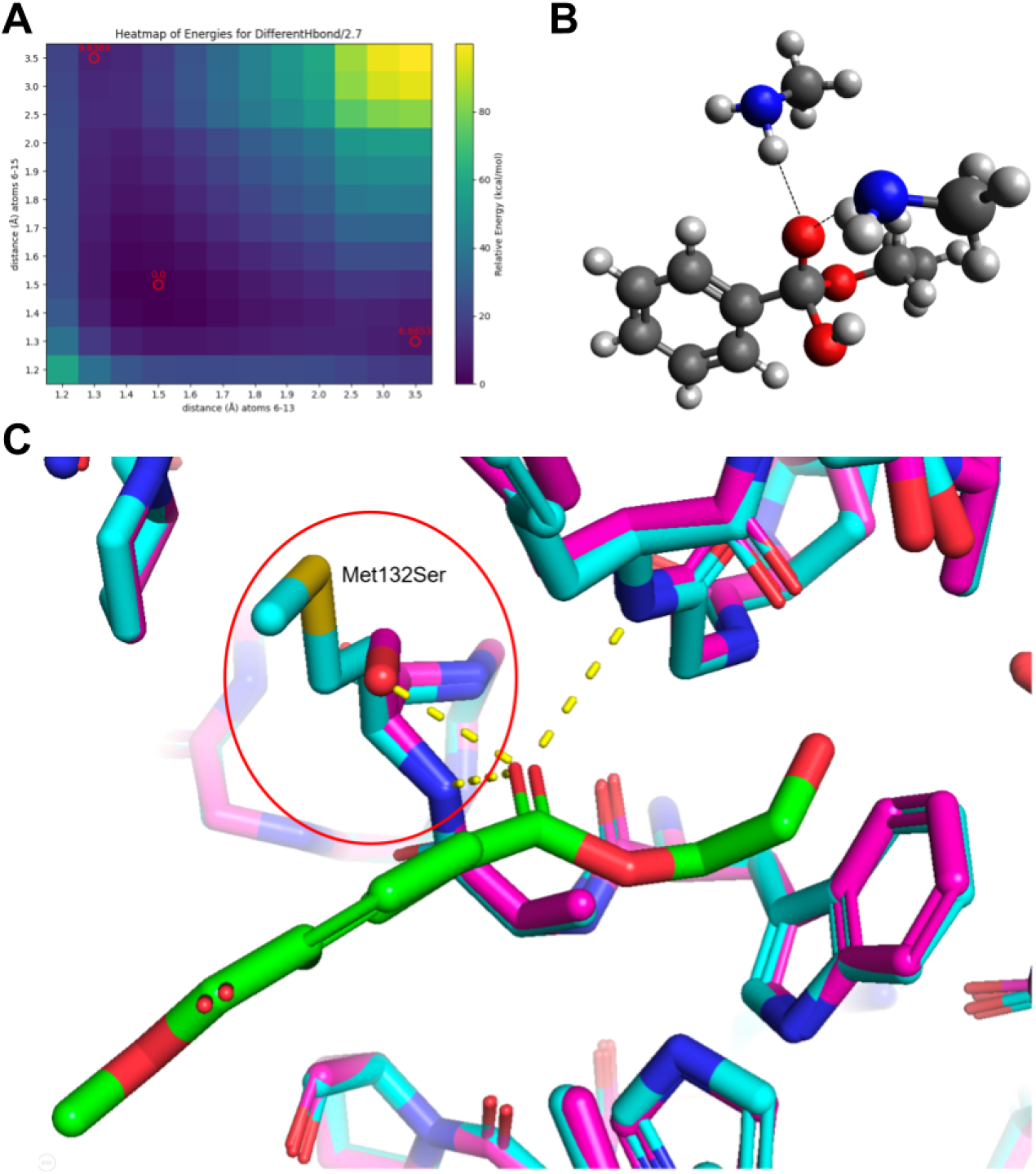
(A) A 2D energy surface scan to locate the transition/intermediate state of the ester hydrolysis reaction with the N(H)-O(=C) distance locked at 2.7Å. X-axis corresponds to the distance between the center C and O(H); Y-axis corresponds to the distance between the center C and O(Et). Thus, the red circle on the bottom right corner represents the reactant; the one on the top left corner represents the product; the red circle in the center represents the transition/intermediate state. (B) Our computational model of the four coordinated transition/intermediate state, with two hydrogen atoms toward the carbonyl. (C) Active site of the AlphaFold 3 predicted model (magenta) overlays with the crystal structure (cyan, PDB: 5XH3). It shows the Met132Ser mutation can form an additional hydrogen bond with the substrate carbonyl.

Thus, we propose mutating the two amides to those with a lower pKa can lower the reaction barrier and thus increase the reaction rate. Interestingly, in IsPETase, the two hydrogen bond donors are Tyr58 and Met132, and their corresponding pKa values are already on the lowest end among all amino acids. Our theory may help to explain why nature chose these two amino acids in the wild type. The only amino acid that has a lower pKa on the amide is Asn, which is a potential candidate to mutate into. On the other hand, we also propose to mutate Met132 to either Ser or Thr, so the OH group on the side chain can form an additional hydrogen bond with the PET carbonyl. We have used AlphaFold 3 to confirm that such mutations do not change the overall folding of the enzyme, and the OH group of a Met132Ser mutation is within the hydrogen bond range with the PET carbonyl.

### 3.5. Big picture of the ester hydrolysis by PETase

The hydrolysis of PET at the PETase active site is a two-step reaction. The first step is an ester exchange reaction by linking the PET to the protein through the key serine, while the follow-up step is to hydrolyze the newly formed ester and regenerate the active site. The two steps are similar in mechanisms, and a general energy profile is plotted in Figure 5.

**Figure 5.**
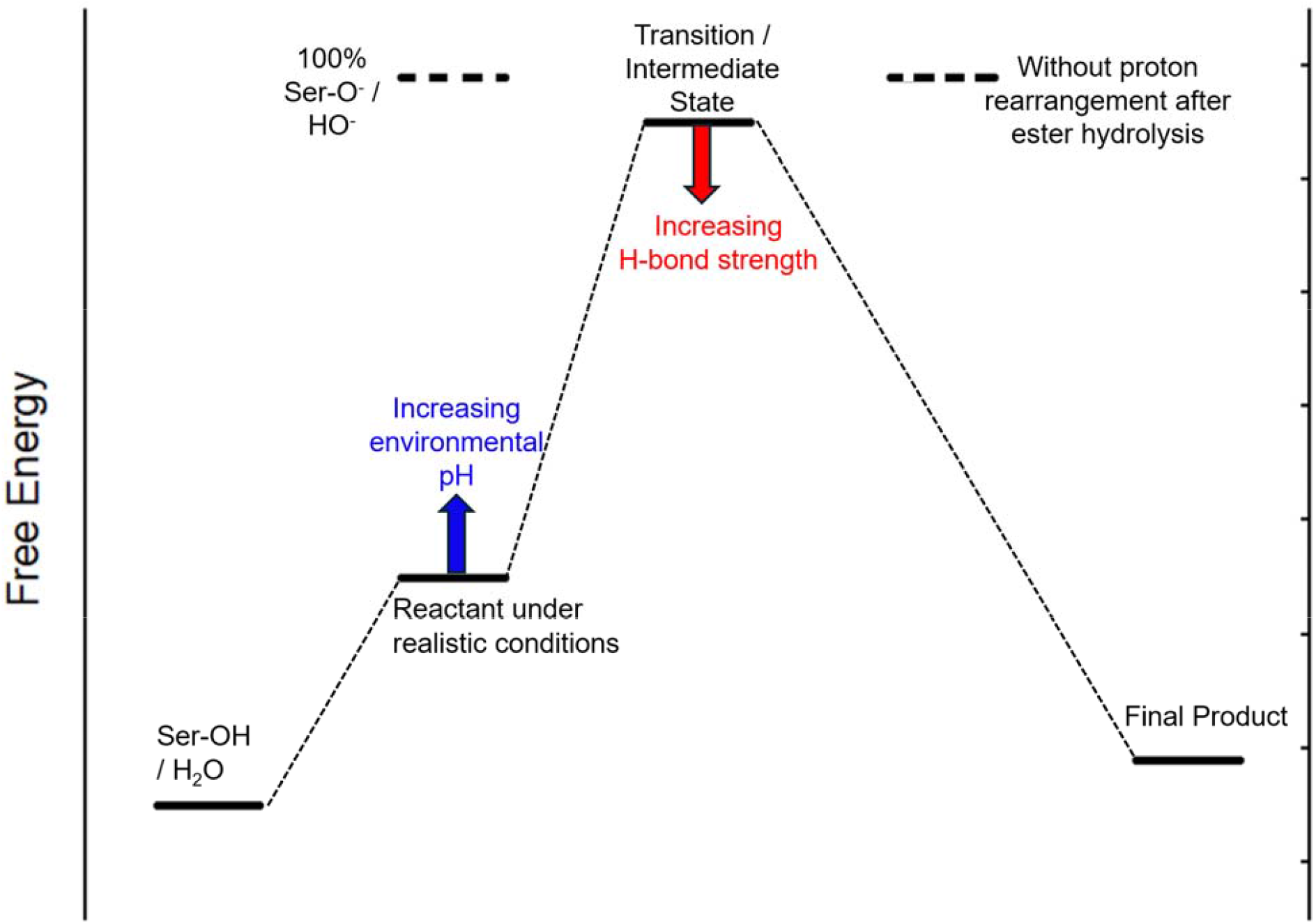
Overall picture of the thermodynamics of the ester hydrolysis. It is demonstrated that increasing the environmental pH can make better reactants, while increasing the H-bond strength towards the PET carbonyl can stabilize the transition/intermediate state. Both ways can improve the efficiency of the enzyme.

It is found that all the C-O bonds in different esters and in the carboxylic acid product have comparable bond strength. The pKa values of water, the OH group in the serine side chain, and the OH group in the alcohol products are also similar, so the reaction is really driven by reactant concentrations. If the hydroxide ion concentration in the environment is increased, there will be a better chance that the serine anion (in step 1) or the hydroxide (in step 2) nucleophile is going to form, so the overall free energy of the reactant is increasing toward the molecular scenario as we calculated for Rxn 1B. On the other hand, the transition/intermediate state can be stabilized by increasing the local hydrogen bond strength. This can be achieved by either mutating the existing amino acids to those that have lower amide pKa’s, such as Asn, or introducing new hydrogen bonds from side chains, such as a Met132Ser mutation.

To sum up, our study brings a fundamental understanding of ester hydrolysis reaction, thus PET degradation reactions themselves at the atomic level. It is shown that the enzyme active site creates a local environment such that it takes advantage of both acidic and basic conditions for the ester hydrolysis. Two directions to engineer the enzyme are then proposed to have better reactivity, either thermodynamically or kinetically. This study also provides novel insights into future research directions and can be extended to other similar enzymes in the family that utilize the oxyanion hole as an active site.

## Acknowledgement

This research used the Lawrencium computational cluster resource provided by the IT Division at the Lawrence Berkeley National Laboratory (Supported by the Director, Office of Science, Office of Basic Energy Sciences, of the U.S. Department of Energy under Contract No. DE-AC02-05CH11231). Colin Zhang and Robert Chuang were supported by the Experiences in Research (EinR) program at the Lawrence Berkeley National Laboratory. Joshua Franklin was supported by the Community College Internship (CCI) program by the Department of Energy.

## Notes

### Competing Interest Statement

The authors have declared no competing interest.

## Reference

(1) Walker, T. R.; Fequet, L. Current Trends of Unsustainable Plastic Production and Micro(Nano)Plastic Pollution. TrAC Trends Anal. Chem. 2023, 160, 116984. 10.1016/j.trac.2023.116984.

(2) Geyer, R.; Jambeck, J. R.; Law, K. L. Production, Use, and Fate of All Plastics Ever Made. Sci. Adv. 2017, 3 (7), e1700782. 10.1126/sciadv.1700782.

(3) Barnes, D. K. A.; Galgani, F.; Thompson, R. C.; Barlaz, M. Accumulation and Fragmentation of Plastic Debris in Global Environments. Philos. Trans. R. Soc. Lond. B. Biol. Sci. 2009, 364 (1526), 1985–1998. 10.1098/rstb.2008.0205.

(4) Andrady, A. L.; Barnes, P. W.; Bornman, J. F.; Gouin, T.; Madronich, S.; White, C. C.; Zepp, R. G.; Jansen, M. A. K. Oxidation and Fragmentation of Plastics in a Changing Environment; from UV-Radiation to Biological Degradation. Sci. Total Environ. 2022, 851, 158022. 10.1016/j.scitotenv.2022.158022.

(5) Wang, J.; Liu, X.; Li, Y.; Powell, T.; Wang, X.; Wang, G.; Zhang, P. Microplastics as Contaminants in the Soil Environment: A Mini-Review. Sci. Total Environ. 2019, 691, 848–857. 10.1016/j.scitotenv.2019.07.209.

(6) Chartres, N.; Cooper, C. B.; Bland, G.; Pelch, K. E.; Gandhi, S. A.; BakenRa, A.; Woodruff, T. J. Effects of Microplastic Exposure on Human Digestive, Reproductive, and Respiratory Health: A Rapid Systematic Review. Environ. Sci. Technol. 2024, 58 (52), 22843–22864. 10.1021/acs.est.3c09524.

(7) The plastic brain: neurotoxicity of micro- and nanoplastics | Particle and Fibre Toxicology. https://link.springer.com/article/10.1186/s12989-020-00358-y (accessed 2025-08-28).

(8) Fan, J.; Ha, Y. Micro- and Nanoplastics and the Immune System: Mechanistic Insights and Future Directions. Immuno 2025, 5 (4), 52. 10.3390/immuno5040052.

(9) Yoshida, S.; Hiraga, K.; Taniguchi, I.; Oda, K. Ideonella Sakaiensis, PETase, and MHETase: From Identification of Microbial PET Degradation to Enzyme Characterization. In Methods in Enzymology; Weber, G., Bornscheuer, U. T., Wei, R., Eds.; Enzymatic Plastic Degradation; Academic Press, 2021; Vol. 648, pp 187–205. 10.1016/bs.mie.2020.12.007.

(10) Han, X.; Liu, W.; Huang, J.-W.; Ma, J.; Zheng, Y.; Ko, T.-P.; Xu, L.; Cheng, Y.-S.; Chen, C.-C.; Guo, R.-T. Structural Insight into Catalytic Mechanism of PET Hydrolase. Nat. Commun. 2017, 8 (1), 2106. 10.1038/s41467-017-02255-z.

(11) Joo, S.; Cho, I. J.; Seo, H.; Son, H. F.; Sagong, H.-Y.; Shin, T. J.; Choi, S. Y.; Lee, S. Y.; Kim, K.-J. Structural Insight into Molecular Mechanism of Poly(Ethylene Terephthalate) Degradation. Nat. Commun. 2018, 9 (1), 382. 10.1038/s41467-018-02881-1.

(12) Lu, H.; Diaz, D. J.; Czarnecki, N. J.; Zhu, C.; Kim, W.; Shroff, R.; Acosta, D. J.; Alexander, B. R.; Cole, H. O.; Zhang, Y.; Lynd, N. A.; Ellington, A. D.; Alper, H. S. Machine Learning-Aided Engineering of Hydrolases for PET Depolymerization. Nature 2022, 604 (7907), 662–667. 10.1038/s41586-022-04599-z.

(13) Stevensen, J.; Janatunaim, R. Z.; Ratnaputri, A. H.; Aldafa, S. H.; Rudjito, R. R.; Saputro, D. H.; Suhandono, S.; Putri, R. M.; Aditama, R.; Fibriani, A. Thermostability and Activity Improvements of PETase from Ideonella Sakaiensis. ACS Omega 2025, 10 (7), 6385–6395. 10.1021/acsomega.4c05142.

(14) Thermostability enhancement of polyethylene terephthalate degrading PETase using self- and nonself-ligating protein scaffolding approaches - Sana - 2023 - Biotechnology and Bioengineering - Wiley Online Library. https://analyticalsciencejournals.onlinelibrary.wiley.com/doi/full/10.1002/bit.28523 (accessed 2025-08-28).

(15) Jerves, C.; Neves, R. P. P.; Ramos, M. J.; da Silva, S.; Fernandes, P. A. Reaction Mechanism of the PET Degrading Enzyme PETase Studied with DFT/MM Molecular Dynamics Simulations. ACS Catal. 2021, 11 (18), 11626–11638. 10.1021/acscatal.1c03700.

(16) Schrödinger, LLC. The PyMOL Molecular Graphics System, Version 3.1, 2015.

(17) Hanwell, M. D.; Curtis, D. E.; Lonie, D. C.; Vandermeersch, T.; Zurek, E.; Hutchison, G. R. Avogadro: An Advanced Semantic Chemical Editor, Visualization, and Analysis Platform. J. Cheminformatics 2012, 4 (1), 17. 10.1186/1758-2946-4-17.

(18) Neese, F. The ORCA Program System. WIREs Comput. Mol. Sci. 2012, 2 (1), 73–78. 10.1002/wcms.81.

(19) Abramson, J.; Adler, J.; Dunger, J.; Evans, R.; Green, T.; Pritzel, A.; Ronneberger, O.; Willmore, L.; Ballard, A. J.; Bambrick, J.; Bodenstein, S. W.; Evans, D. A.; Hung, C.-C.; O’Neill, M.; Reiman, D.; Tunyasuvunakool, K.; Wu, Z.; Žemgulytė, A.; Arvaniti, E.; Beattie, C.; Bertolli, O.; Bridgland, A.; Cherepanov, A.; Congreve, M.; Cowen-Rivers, A. I.; Cowie, A.; Figurnov, M.; Fuchs, F. B.; Gladman, H.; Jain, R.; Khan, Y. A.; Low, C. M. R.; Perlin, K.; Potapenko, A.; Savy, P.; Singh, S.; Stecula, A.; Thillaisundaram, A.; Tong, C.; Yakneen, S.; Zhong, E. D.; Zielinski, M.; Žídek, A.; Bapst, V.; Kohli, P.; Jaderberg, M.; Hassabis, D.; Jumper, J. M. Accurate Structure Prediction of Biomolecular Interactions with AlphaFold 3. Nature 2024, 630 (8016), 493–500. 10.1038/s41586-024-07487-w.

(20) Xu, T.; Chen, J.; Wang, Z.; Tang, W.; Xia, D.; Fu, Z.; Xie, H. Development of Prediction Models on Base-Catalyzed Hydrolysis Kinetics of Phthalate Esters with Density Functional Theory Calculation. Environ. Sci. Technol. 2019, 53 (10), 5828–5837. 10.1021/acs.est.9b00574.

(21) Functional and Structural Characterization of PETase SM14 from Marine-Sponge Streptomyces sp. Active on Polyethylene Terephthalate | ACS Sustainable Chemistry & Engineering. https://pubs.acs.org/doi/10.1021/acssuschemeng.5c00737 (accessed 2025-11-05).

(22) Šebesta, F.; Sovová, Ž.; Burda, J. V. Determination of Amino Acids’ pKa: Importance of Cavity Scaling within Implicit Solvation Models and Choice of DFT Functionals. J. Phys. Chem. B 2024, 128 (7), 1627–1637. 10.1021/acs.jpcb.3c07007.

